# The scorpionfly (*Panorpa cognata*) genome highlights conserved and derived features of the peculiar dipteran X chromosome

**DOI:** 10.1101/2023.07.11.548499

**Authors:** Clementine Lasne, Marwan Elkrewi, Melissa A. Toups, Lorena Layana, Ariana Macon, Beatriz Vicoso

## Abstract

Many insects carry an ancient X chromosome - the Drosophila Muller element F - that likely predates their origin. Interestingly, the X has undergone turnover in multiple fly species (Diptera) after being conserved for more than 450 MY. The long evolutionary distance between Diptera and other sequenced insect clades makes it difficult to infer what could have contributed to this sudden increase in rate of turnover. Here, we produce the first genome and transcriptome of a long overlooked sister-order to Diptera: Mecoptera. We compare the scorpionfly *Panorpa cognata* X-chromosome gene content, expression, and structure, to that of several dipteran species as well as more distantly-related insect orders (Orthoptera and Blattodea). We find high conservation of gene content between the mecopteran X and the dipteran Muller F element, as well as several shared biological features, such as the presence of dosage compensation and a low amount of genetic diversity, consistent with a low recombination rate. However, the two homologous X chromosomes differ strikingly in their size and number of genes they carry. Our results therefore support a common ancestry of the mecopteran and ancestral dipteran X chromosomes, and suggest that Muller element F shrank in size and gene content after the split of Diptera and Mecoptera, which may have contributed to its turnover in dipteran insects.

## Introduction

Sex chromosomes originally arise from autosomes (Muller 1914; Ohno 1967), but over time can evolve highly specialized sequence and regulatory features. Loss of recombination between nascent X and Y chromosomes often leads to genetic degeneration of the Y, which becomes gene-poor and enriched for transposable elements and other repeats (Charlesworth et al. 1994). This degeneration can cause gene expression imbalances between X-linked and autosomal genes in the heterogametic sex, which in turn select for the evolution of dosage compensation mechanisms that re-establish optimal X:autosomes expression balance, such as doubling the expression of the male X in *D. melanogaster* (Gupta et al. 2006). Finally, insect X chromosomes are often enriched for genes that are primarily expressed in females (female-biased genes), and depleted of male-biased genes (Parisi et al. 2003; Mikhaylova and Nurminsky 2011; Pal and Vicoso 2015; Whittle et al. 2020; Parker et al. 2022). Due to these unusual features, highly differentiated sex chromosomes are thought to be difficult to revert to autosomes and to be maintained over long periods of time, or even become non-reversible “evolutionary traps” (Pokorná and Kratochvíl 2009). The growing pool of genomic and transcriptomic data for both model and non-model organisms has provided support for the long-term existence of stable sex chromosomes with highly conserved gene content across entire clades - such as the XY chromosomes of eutherian mammals and the avian ZW chromosomes (Marshall Graves 2016; Vicoso 2019), but also uncovered clades with high rates of sex-chromosome turnovers between closely related species, e.g. frogs (Jeffries et al. 2018), cichlids (El Taher et al. 2021) and crustaceans (Becking et al. 2017). It remains unclear why some taxa acquire highly conserved sex chromosomes and others have very high rates of turnover.

Insects are an excellent taxon to study both conservation and turnover of sex chromosomes. They show both male- and female-heterogametic systems, as well as tremendous variation in the extent of sex-chromosome (and gene content) conservation between different orders (Blackmon et al. 2017). Recurrent sex-chromosome turnover has occurred in flies (Diptera), where the ancestral sex chromosome (the dipteran “Muller element F”) has been replaced as the X by another chromosome multiple times independently (Vicoso and Bachtrog 2013; Vicoso and Bachtrog 2015). On the other hand, conservation of the X chromosome has been observed in Hemipterans (Pal and Vicoso 2015) and Coleoptera (Bracewell et al. 2023). The most striking evidence of conservation so far is the apparent homology between the X chromosomes of the cockroach (Blattodea) (Meisel et al. 2019), the damselfly (Chauhan et al. 2021), the grasshopper (Orthoptera) (Li et al. 2022) and the ancestral dipteran X chromosome - element F, suggesting that the same X chromosome has been maintained for over 400 millions years of evolution. Why such an ancient and well conserved sex chromosome would undergo repeated turnover in Diptera is unclear, but may have to do with its reduced size and gene content in this clade, which should mitigate the fitness consequences of reverting it to an autosome (Vicoso 2019; Toups and Vicoso 2023). However, due to the very large evolutionary distances between these insects, it is difficult to conclusively disentangle whether there is long-standing conservation of Muller element F as the X chromosome across insect orders, or if this represents the convergent recruitment of the same set of genes for sex determination. Element F is also known to have an unusual biology in *Drosophila melanogaster*, where it has been studied extensively: it is almost entirely heterochromatic and does not undergo crossing over (and consequently has an extremely low recombination rate). Whether these features are related to its small size and/or turnover as the X chromosome is unclear, since no close outgroup of Diptera carrying element F as the X has been characterized.

Relatively few molecular and genomic resources are available for Mecoptera - the sister-order to Diptera that comprises scorpionflies and hangingflies (Misof et al. 2014). Cytogenetic studies show that almost all Mecoptera species studied so far are XX/XO (Miao et al. 2019), but the mecopteran X chromosome has not yet been characterised at the molecular level. Intriguingly, it has been described as “dot-shaped” in meiotic spreads of several *Panorpa* scorpionfly species (Xu et al. 2013), a term that is reminiscent of the shape of Muller element F in Drosophila (where it is also known as the “dot chromosome” (Ashburner et al. 2005)), making scorpionflies a promising model for understanding the evolution of the peculiar element F. We produced a high-quality genome assembly from PacBio reads for the scorpionfly species *Panorpa cognata* (order: Mecoptera). We identified X-derived scaffolds, and inferred the level of conservation of gene content of the X chromosome between this clade and various dipteran and non-dipteran insects. We combined our genome assembly with extensive transcriptomic data to explore patterns of dosage compensation in different tissues and tissue-specificity of X and autosomal genes. Finally, we investigated whether the *P. cognata* X showed features of a heterochromatic chromosome, similar to Muller element F.

## Methods

### Sample collection and sequencing

*P. cognata* specimens were collected in August 2021 in Maria Gugging (Lower Austria) and immediately frozen at −80°C until further processing. Species identification was confirmed by sequencing the mitochondrial cytochrome c oxidase I (COI) gene and comparing it to available sequences for this species (Misof et al. 2000). High molecular weight DNA was extracted from a single male with the Qiagen Genomic-Tip 100/G Kit, and used for PacBio long read DNA sequencing. A single frozen female was used for Hi-C library prep and illumina sequencing. For illumina whole genome sequencing, DNA was extracted from 1 male and 1 female separately using the Qiagen DNeasy Blood and Tissue kit and fragmented using the Bioruptor Plus Ultrasonicator. Total RNA was extracted from the heads, gonads and carcasses of the 3 males and 3 females (samples were not pooled) using the Bioline Isolate II RNA extraction kit, resulting in 3 biological replicate samples per tissue and sex and a total of 18 libraries. All DNA and RNA sequencing libraries were prepared and sequenced at the Vienna Biocenter Sequencing Facility. All RNA and DNA samples used for the downstream transcriptome assembly and gene expression analysis are listed in **Table S1**, and the corresponding sequencing reads are available at the NCBI Short Reads Archive under Bioproject number PRJNA989034.

### Genome assembly

PacBio consensus reads were generated from the raw bam file using the PacBio CCS tool (version 6.4.0, on conda 4.14.0). The CCS reads were assembled using Hifiasm (version 0.15-r327; (Cheng et al. 2021)), and the primary assembly was purged using purge_dups (version 1.2.5; (Guan et al. 2020)) to remove any duplicate sequences. The Hi-C reads were then aligned to contigs longer than the N80 of the assembly (as smaller contigs still appeared to be largely redundant), and processed using the HiC-Pro pipeline (version 3.1.0; (Servant et al. 2015)). The valid alignments were extracted from the resulting bam file, further filtered for edit distance (NM:i:0) using Matlock (phase genomics), and then used for scaffolding the purged primary assembly with YaHS (YaHS-1.2a.1.patch; (Zhou et al. 2023)). BUSCO was used to assess the completeness of the genome with the arthropoda_odb10 dataset (version 5.4.4; (Manni et al. 2021)). As most of the genome is contained in super-scaffolds, we performed downstream analyses using the longest 25 scaffolds (**Table S2**). The choice of scaffold number was mainly based on the large drop in length after the 25th scaffold and supported by the minor decrease in BUSCO score (**Figure S1**).

### Identification of X-linked scaffolds

The *P. cognata* female and male Illumina DNA reads were mapped to the assembled genome using Bowtie2 (version 2/2.4.5; (Langmead and Salzberg 2012)) with end-to-end sensitive mode. SOAP.coverage (version 2.7.7; https://github.com/gigascience/bgi-soap2/tree/mas-ter/tools/soap.coverage) was used to calculate the genomic coverage for each scaffold in windows of 10000 bp from the resulting SAM alignment. The log_2_ of the ratio of male to female coverage was calculated for all the windows and the [median(log_2_(Male/Female coverage))-0.5] was used as a cut-off to assign scaffolds as either X-linked or autosomal. If the median(log_2_(Male/Female coverage)) for the scaffold windows was below the cut-off, the scaffold was assigned as X-linked, otherwise it was assigned as an autosome.

### Transcriptome assembly and transcripts genomic location

The *P. cognata* transcriptome was assembled from all 18 RNA-seq libraries. Quality control of the paired-end reads was conducted using FastQC (version 0.11.9; (Andrews 2010)) and quality filtering with TRIMMOMATIC (version 0.36; (Bolger et al. 2014)). We used Trinity (trinityrnaseq-v2.11.0; (Grabherr et al. 2011)) and Evigene (EvidentialGene tr2aacds.pl version 2022.01.20; (Gilbert 2016)) to assemble and curate the transcriptome, and further filtered for transcript sequence-length greater than 500bp using fafilter (UCSC source code collection, http://genome.ucsc.edu/). The transcriptome assembly quality was checked with BUSCO using arthropoda_odb10 as a reference dataset (version 5.4.4; (Manni et al. 2021)). The final transcriptome assembly consists of 36618 transcripts and is available at the ISTA data repository *[a permanent URL will be added upon acceptance]*.

To determine the genomic location of each transcript, we mapped our transcriptome to our genome assembly with Standalone BLAT (version 36×2; (Kent 2002)). We used custom Perl scripts to keep only the best hit for each gene in the genome and, when multiple transcripts overlapped on the genome, to keep only the transcript with the highest mapping score (unless they overlapped by less than 20 bps, in which case both were kept).

### Homology of the Panorpa cognata and Cochliomyia hominivorax X chromosomes

The *P. cognata* protein sequences were obtained from the transcriptome using a Perl script (GetLongestAA_v1_July2020.pl), which outputs the longest amino acid sequence for each *P. cognata* transcript. The published annotation file (GFF) and genome of the New World screwworm *Cochliomyia hominivorax* (order: Diptera; suborder: Brachycera) were obtained from Dryad (Scott 2022). The protein sequences of *C. hominivorax* were extracted from the GFF and genome files using gffread (version 0.12.7; (Pertea and Pertea 2020)) and were filtered with a Perl script (GetLongestCDS_v2.pl) to get the longest isoform per protein. The correspondence between Muller elements and *C. hominivorax* chromosomes was obtained from Tandonnet et al. (2023). Since an outgroup was required to obtain orthologous genes between the two species, the protein sequences of the yellow fever mosquito *Aedes aegypti* (order: Diptera; suborder: Nematocera) were retrieved from Ensembl Metazoa (and were also filtered with the Perl script mentioned above).

We then used Orthofinder (Emms and Kelly 2019) to obtain 1-to-1 orthologous genes between *C. hominivorax* and *P. cognata*, and calculated the proportion of these 1-to-1 orthologs that were X-linked in *P. cognata* (hereafter “X-linkage threshold”). We then estimated the proportion of *P. cognata* genes that are X-linked separately for 1-to-1 orthologs that are on each Muller element of *C. hominivorax*. We performed a chi-squared test comparing the proportion obtained for each Muller element to the proportion obtained from all the others (e.g. element A versus elements B,C,D,E,F), using the Python function scipy.stats.chi2_contingency from SciPy library (Virtanen et al. 2020). Muller elements that had a significant p-value (*P* < 0.05) and were above “the X-linkage threshold” were considered as overrepresented. We also performed this analysis using *Drosophila melanogaster* (**Methods S1**).

### Conservation of X-linked gene content between P. cognata and other insects

We assessed whether the X-linked genes of *P. cognata* were also present on the X chromosome of two other insect species: the migratory locust *Locusta migratoria* (Order: Orthoptera) and the spotted crane fly *Nephrotoma appendiculata* (suborder: Nematocera, a basal dipteran known to have element F as the X). The tree representing the phylogenetic relationship between these species was generated using the online tool iTol (https://itol.embl.de/about.cgi, version 6.7.3) based on the topology of Misof et al. (2014). Since a genome annotation was not available for these species we used a pipeline that bypassed the need for protein sequences to infer homology between X chromosomes. We downloaded chromosome-level genome assemblies from the National Center for Biotechnology Information (NCBI) for *L. migratoria (*https://www.ncbi.nlm.nih.gov/assembly/GCA_026315105.1/) and *N. appendiculata (*https://www.ncbi.nlm.nih.gov/assembly/GCA_947310385.1. We then used Standalone BLAT (version 36×2; (Kent 2002)) to map our *P. cognata* transcriptome to the genome of these two species using a translated query and database, and filtered for hits with a match score above 50. A Perl script (1-besthitblat.pl) was then used to get only the best hit for each transcript in the genome, and another Perl script (2-redremov_blat_V2.pl) to keep only the transcript with the highest mapping score when two transcripts overlapped by more then 20bps. We used this set of *P. cognata* transcripts, with their genomic location in *L. migratoria* and *N. appendiculata*, as a proxy for the location of orthologous genes.

### Synteny of P. cognata, C. hominivorax and N. appendiculata

Synteny was examined between *P. cognata* and two dipteran species, *C. hominivorax* and *N. appendiculata*, using GENESPACE (version 0.94; (Lovell et al. 2022)), which requires a GFF annotation and a set of peptide sequences for each species. For *C. hominivorax*, we used the GFF provided by the *C. hominivorax* genome project and the peptide sequences produced as described above as input. For the other species, new amino acid sequences that met the GENESPACE input requirements were obtained. We obtained a genome annotation for *P. cognata* by mapping the RNA-seq libraries to the genome using HISAT2 (version 2.2; (Kim et al. 2019)). GTF files were generated for each library and then merged together using StringTie2 (version 2.2.1; (Kovaka et al. 2019)). The resulting GTF file was then converted to the GFF3 format using the gffread command from the cufflinks package (cufflinks version 2.2.1; (Trapnell et al. 2010)). We input the StringTie GTF file produced above into Transdecoder (version 5.5; Haas, BJ. https://github.com/TransDecoder/TransDecoder)) to select the longest ORFs. We then searched for homology between our ORFs and the uniprot database (The UniProt Consortium et al. 2023) using ncbi blast (version 2.2.31; (Camacho et al. 2009)). Blast results were integrated into Transdecoder (version 5.5.0; Haas, BJ. https://github.com/TransDecoder/TransDecoder) protein prediction. We then selected the longest isoform using a custom Perl script.

To generate peptide sequences for *N. appendiculata* and *L. migratoria*, we first downloaded RNAseq for each species (ERR10378025 (https://www.ncbi.nlm.nih.gov/sra/?term=-ERR10378025) and SRR22110765 (Li et al. 2022), respectively) from the Sequence Read Archive hosted by NCBI. Quality was assessed using FastQC (https://www.bioinformatics.babraham.ac.uk/projects/fastqc/). Reads were quality trimmed and adapters were removed with Trimmomatic (version 0.39; (Bolger et al. 2014)). We then proceeded with the pipeline described in the previous paragraph for *P. cognata* to produce a GFF file and peptide sequences for the longest isoform of each gene.

### Gene expression and dosage compensation

#### Quantification and normalisation

The newly assembled *P. congata* transcriptome was indexed with Kallisto (version 0.46.2; (Bray et al. 2016)). The trimmed RNA-seq reads of all 18 samples were mapped to the transcriptome and gene expression was quantified using the same program. Only transcripts mapping to the largest 25 scaffolds in the genome were retained for further analyses. Further gene expression and statistical analyses were performed in R (R Core Team 2020). We performed quantile normalisation of gene expression (in Transcripts Per Million, TPM) across all 18 samples using the R package NormalizerDE (version 1.16.0; (Willforss et al. 2019)). We then visualised the overall similarity in expression profiles of our samples using the Spearman correlation option embedded in the function heatmap.2 of the R package gplots (version 3.1.3; https://github.com/talgalili/gplots).

#### Dosage compensation

For each tissue, gene expression was first normalised across male and female samples, then averaged within each sex. A second quantile normalisation was applied to these sex averages, and only genes with expression levels > 0.5 TPM in both sexes were kept for comparing expression patterns between the X and autosomes. Significant differences in gene expression values between sexes and chromosomes were tested for using Wilcoxon rank sum tests.

#### Tissue-specific expression

Tissue-specific expression of autosomal and X-linked genes was assessed by averaging gene expression across both sexes for heads and for carcasses, but separately for gonads to obtain testis-specific and ovary-specific gene expression. A gene was considered as tissue-specific if its expression level was greater than 1 TPM in a tissue and smaller than 0.5 TPM in all other tissues. Significant differences in the proportions of tissue-specific genes between the X and the autosomes were assessed using the chi-squared test option in the pairwiseNominalIndependence function of the R package rcompanion (version 2.4.21; (Mangiafico 2023)).

#### Sex-biased gene expression

Genes that are differentially expressed between the two sexes in gonads, heads, and carcasses were called using the R package sleuth (Pimentel et al. 2017). Genes with q-values < 0.05, a TPM value > 0.5 in both sexes and a 2-fold differential expression between the sexes were considered sex-biased. Significant differences in the proportions of sex-biased genes between the X and the autosomes were assessed using the chi-squared test option in the pairwiseNominalIndependence function of the R package rcompanion. Statistical analyses could not be conducted in head and carcass, as too few genes were sex-biased in these tissues.

### GC content and nucleotide diversity

GC content was estimated for 10000 bp windows along the genome scaffolds using the GCcalc.py script (https://github.com/WenchaoLin/GCcalc). To assess the nucleotide diversity of the transcriptome, the RNAseq reads were first aligned to the transcriptome using bwa-mem (Li 2013), and then SNPs were called using bcftools (Danecek et al. 2021) and filtered using vcftools (Danecek et al. 2011). The filtered vcf file was then used as input to PIXY (Korunes and Samuk 2021), which calculates the population genetic summary statistic pi (ν), with a sliding window size of 28kb (corresponding to the largest transcript in our data, such that we obtained one value of pi per transcript).

### Repeat Content

A consensus repeat library was generated and annotated using RepeatModeler (version 2.0.4; (Flynn et al. 2020)). The repeat library was used with RepeatMasker (version 4.1.5; (Smit et al. 2013)) to get a detailed annotation of the repeat content across the genome. The proportion of repeats were obtained for windows of 10000 bp from the output of RepeatMasker using a custom Python script.

## Results

### Genome assembly and identification of the X

We produced the first mecopteran genome assembly for the species *P. cognata*, using PacBio reads from a single male and illumina Hi-C reads from a single female. The final genome assembly contains 187 scaffolds, and the estimated genome size is 0.46 GB. The BUSCO analysis revealed a 99% genome assembly completeness (**Figure S1**, left panel). Although the assembly is not chromosome-level (cytogenetic studies of *P. cognata* reported n=22 chromosomes (Miao et al. 2019)), potentially due to the low complexity/quality of the Hi-C data, most of the genome is contained in super-scaffolds. In particular, 73% of the genome is in the longest 25 scaffolds, which we focus on for the rest of the manuscript (the corresponding BUSCO score is 93%). Based on their reduced ratio of male to female short read genomic coverage, two super-scaffolds (scaffold_1 and scaffold_22) and a few smaller scaffolds were identified as X-linked (**Figure 1**). The absence of scaffolds with male-specific coverage in the genome assembly supports the lack of a Y chromosome in *P. cognata* (**Figure S2**).

**Figure 1:**
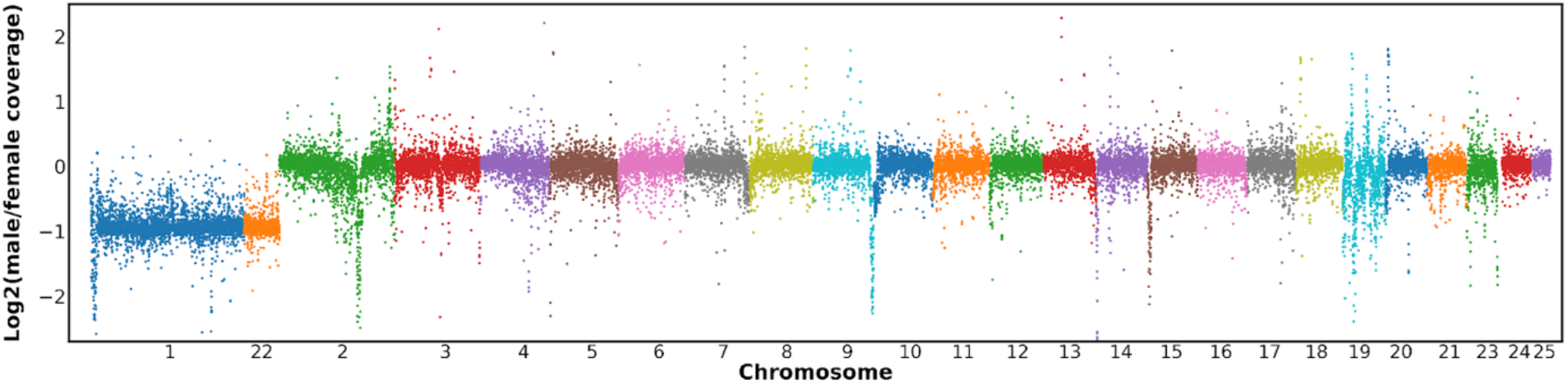
Patterns of male/female coverage for the longest 25 scaffolds in 10000 bp windows. Scaffolds 1 and 22 were classified as X-linked based on their reduced male:female coverage ratio.

### Conservation of the X chromosome

To identify X-linked genes, we assembled a transcriptome (see methods and next section), which we mapped to the *P. cognata* genome. Of the 13214 non-redundant mapped transcripts, a proxy for individual genes, 1520 (11.5%) mapped to X-linked scaffolds, showing that the X is one of the largest and most gene-rich chromosomes in this species. We then investigated whether this X chromosome was homologous to the X of several other insects (**Figure 2(a)**).

**Figure 2:**
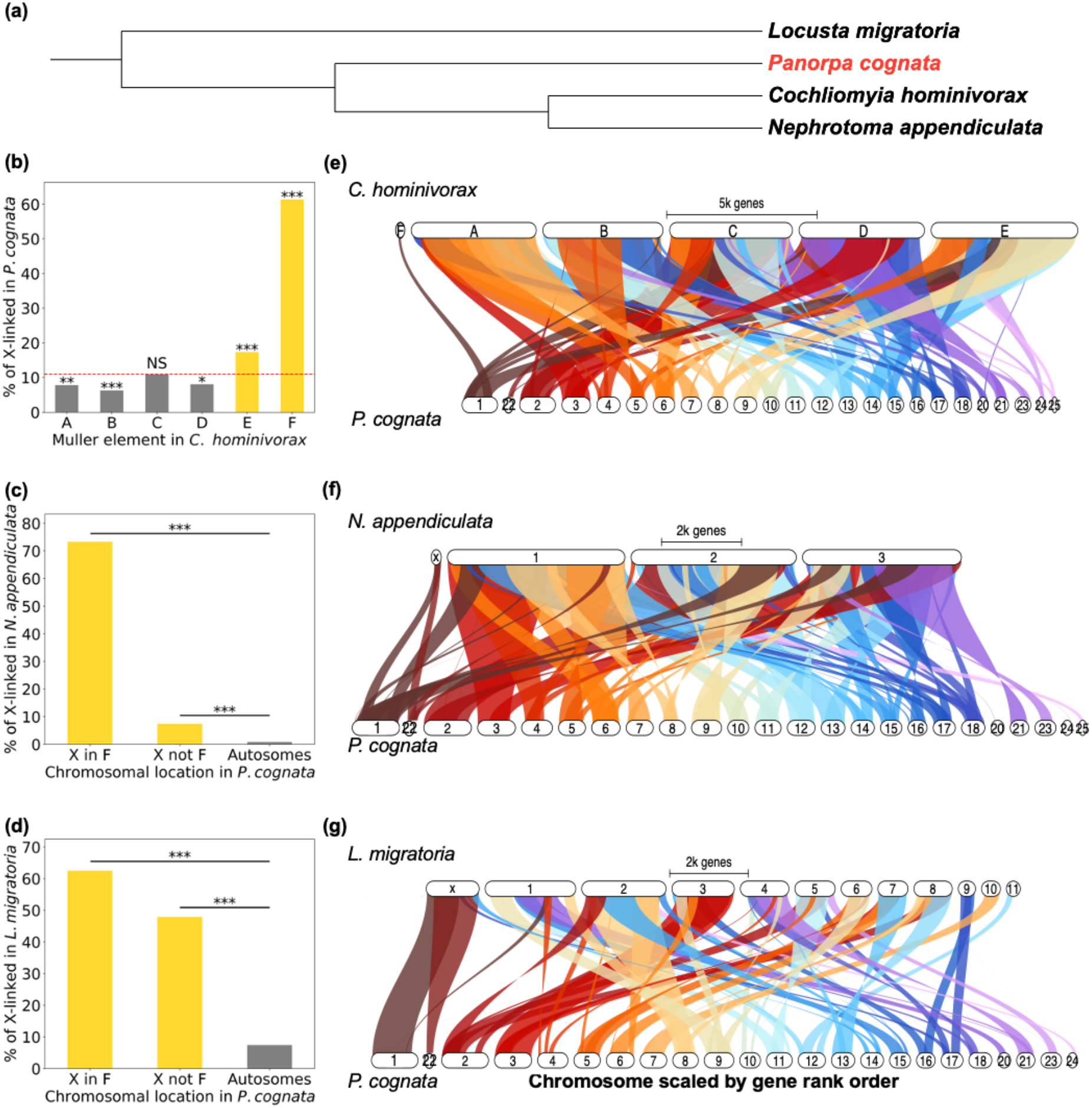
Homology of the X chromosomes of *P. cognata* (order: Mecoptera) and three other insects: two Diptera, *C. hominivorax* (suborder: Brachycera) and *N. appendiculata* (suborder: Nematocera), and *L. migratoria* (order: Orthoptera). (a) Phylogenetic tree of the 4 species. (b) Percentage of genes on each of the *C. hominivorax*’s Muller elements that are X-linked in *P. cognata*. The red dashed line represents the overall proportion of orthologs that are X-linked in *P. cognata* (i.e. the “X-linkage threshold”). (c) Percentage of X-linked and autosomal *P. cognata* genes that are X-linked in *N. appendiculata.* The X-linked genes of *P. cognata* were divided into two sets, based on whether they were F-linked in *C.hominivorax* (X-in-F), or not (X-not-F). (d) same as (c) but showing the percentage of *P. cognata* genes that are X-linked in *L. migratoria.* Statistically significant differences between observed and expected percentages were assessed using a chi-squared test (* *P* < 0.05, ** *P* < 0.01, *** *P* < 0.001, NS not significant). (e,f,g) Synteny plots between *P. cognata*’s 25 largest genome scaffolds and the genomes of the 3 other insect species.

We first tested for homology between the *P. cognata* and dipteran X chromosomes by detecting 1-to-1 orthologs with the screwworm *C. hominivorax*, a dipteran species that has maintained the ancestral element F as the X. **Figure 2(b)** shows that *C. hominivorax* genes located on the X-linked element F, and to a lesser extent on the autosomal element E, are significantly overrepresented among *P. cognata* X-linked genes. The overrepresentation of those elements also holds when taking into account all the scaffolds in our *P. cognata* genome (**Figure S3(a)**) and when *D. melanogaster* is used as the dipteran outgroup (**Figure S4**). The synteny plot between the *C. hominivorax* genome and the 25 largest scaffolds from our *P. cognata* genome (**Figure 2(e)**) supports the homology of the *P. cognata* X-linked scaffolds 1 and 22 to Muller elements E and F in *C. hominivorax*, despite the poor conservation of synteny overall.

The previous results show that, while the *P. cognata* and dipteran X chromosomes are homologous, many *P. cognata* X-linked genes are derived from other Muller elements. We first set out to test if this additional gene content of the X reflects the ancestral state of insects, or instead corresponds to an increase in X-linked gene content in the *P. cognata* lineage. To do so, we divided the *P. cognata* X-linked genes into two based on the location of their homologues in the *C. hominivorax* genome: a set homologous to dipteran element F genes (“X in F”), and a set homologous to genes on other chromosomes (“X not F”). We then estimated the proportion of the two sets that are also X-linked in the distant outgroup *L. migratoria* (and *B. germanica* in **Figure S5**). **Figure 2(d)** shows that the percentage of “X not F” genes that are also X-linked in *L. migratoria* (∼45%) is greater than the corresponding percentage for *P. cognata* autosomal genes (<10%, *P* < 0.001, chi-squared test), suggesting that the difference in gene content reflects at least partly a loss of X-linked genes in dipterans (in agreement with (Toups and Vicoso 2023)). We performed a similar analysis with *N. appendiculata*, a dipteran species that is a putative outgroup to flies and mosquitoes, to investigate if the shrinking of the X occurred early in dipteran evolution, or later in the Brachycera (“higher dipterans”). While there is still an excess of *P. cognata* “X not F” genes on the *N. appendiculata* X (*P* < 0.001, chi-squared test), the percentage (∼10%) is much lower than in the previous analysis with the locust. This suggests that much of the loss of genes in the ancestral X chromosome of Diptera occurred at some point before the split of Tipulidae. We obtained similar results when we considered all the scaffolds in our *P. cognata* genome (**Figure S3(b-c)**).

### Gene expression of the X, dosage compensation, and gene content

The *P. cognata* transcriptome was assembled by pooling male and female head, gonad, and carcass samples (See Methods). The final transcriptome contains 36618 transcripts and has an N50 of 1329 bp. The completeness of our transcriptome assembly was estimated to 92.7% according to our BUSCO analysis (**Figure S1,** right panel). All subsequent analyses of the gene expression data were conducted using the 12357 transcripts of known location on the first 25 genome scaffolds: 11083 on the autosomes and 1274 on the X. A Spearman correlation analysis confirmed that the RNA-seq samples cluster together according to tissues, and according to sex within the gonad and carcass clusters (**Figure S6**).

We compared male and female gene expression on the autosomes and on the X in heads, a somatic organ, and gonads, to assess patterns of dosage compensation and sex-biased expression in scorpionflies (**Figure 3**). We found no difference in expression between autosomal and X-linked genes, nor between the sexes, in heads (**Figure 3(a)**). The male-over-female expression ratio was also similar between autosomal and X-linked genes in this tissue (**Figure 3(c)**), confirming that the X chromosome is fully compensated. In gonads, the expression of X-linked genes was significantly lower in males relative to females (*P* < 0.001, Wilcoxon rank sum test; **Figure 3(b)**) and relative to male autosomal genes (*P* < 0.001, Wilcoxon rank sum test). The male-over-female expression ratio of the X chromosome was also significantly lower than that of autosomes (*P* < 0.001, Wilcoxon rank sum test; **Figure 3(d)**), suggesting that either dosage compensation is incomplete in this tissue, or that a differential accumulation of genes with sex-biased expression has occurred on the X chromosome (see below). Similarly to heads, we found evidence of dosage compensation in carcasses (**Figure S7).**

**Figure 3:**
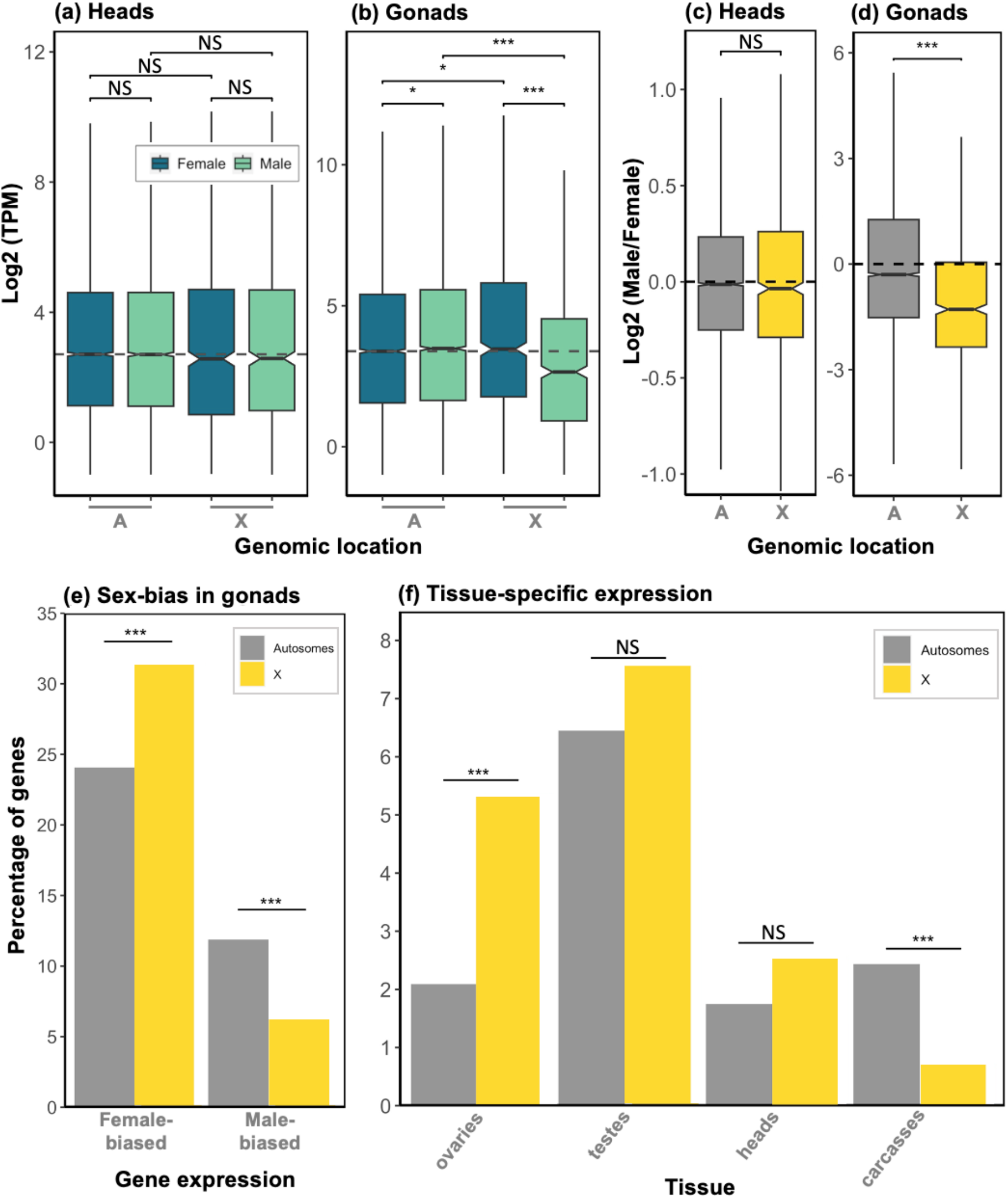
Dosage compensation and biased gene content of the X. (a) and (b): expression of autosomal and X-linked genes in males and females, in heads and gonads, respectively (grey dashed line is the female autosomal gene expression median). (c) and (d): Log_2_ of male-over-female expression ratios for the autosomal and X-linked genes, in heads and gonads, respectively. Statistically significant differences between groups were assessed using a Wilcoxon rank sum test (* adj. *P* < 0.05, ** adj. *P* < 0.01, *** adj. *P* < 0.001, NS not significant). (e) percentage of autosomal and X-linked genes exhibiting sex-biased expression in gonads. (f) Percentage of autosomal and X-linked genes showing tissue-specific expression. Statistically significant differences between the autosomes and the X in (e) and (f) were assessed using a chi-squared test.

The Drosophila gene *Painting-of-fourth* (*POF*) has been shown to mediate dosage compensation in the sheep blowfly, *Lucilia cuprina*, a dipteran species with the ancestral element F as the X. Given the homology between the *P. cognata* and dipteran X chromosomes, we investigated whether *POF* showed patterns of expression consistent with a role in dosage compensation in scorpionflies, i.e. whether it was expressed primarily in male somatic tissues, but less so in testis and in female tissues. Contrary to this, *POF* seemed to be expressed at similar levels heads of both sexes and in ovaries, but showed reduced expression in testes and to some extent in carcasses (**Figure S8**).

Finally, X chromosomes often differ from autosomes in the proportion of sex-biased, tissue- and sex-specific genes that they carry. We found an excess of female-biased genes on the X relative to the autosomes in gonads: 31.6% and 24.1%, respectively (*adj P* < 0.01, chi-squared test; **Figure 3(e)**). We also observed a paucity of male-biased genes on the X (6.1%) relative to the autosomes (11.9%) (*adj P* < 0.001, chi-squared test). Fewer than 50 genes were found to be sex-biased in carcasses, and only one gene in heads, such that no comparisons between the X and the autosomes were possible. We also investigated the extent to which X-linked and autosomal genes show tissue-specific expression. We found a significantly greater proportion of genes with ovary-specific expression on the X chromosome relative to the autosomes: 5.34% and 2.09% of genes, respectively (*adj. P* < 0.001, chi-squared test; **Figure 3(f)**). However, there was no difference in gene-specificity between the X and autosomes in heads and testes, and we note a high percentage of testis-specific genes on the X (7.54%). Finally, the percentage of genes showing carcass-specific expression was significantly lower on the X relative to the autosomes: 0.7% and 2.44%, respectively (*adj. P* < 0.001, chi-squared test).

### X vs. autosomal genetic diversity, CG and repeat content

Muller element F, which corresponds to the ancestral dipteran X, is non-recombining and largely heterochromatic in Drosophila. We investigated whether these features might have already been present in the X of the ancestor of dipterans and mecopterans. We compared autosomal and X-linked pairwise nucleotide diversity (π) and found that autosomal scaffolds have higher levels of genetic diversity than X-linked scaffolds in both sexes (*P* < 0.001, Wilcoxon rank sum test; **Figure 4(a)** for females). The X/Autosome diversity is 0.23 in females, and 0.12 in males, well below the expectation of X/Autosome = 0.75 (the null hypotdhesis when only the number of copies of X chromosomes and autosomes in a population are considered), consistent with a low recombination rate of the X. GC content, which is also correlated with recombination rate (Charlesworth et al. 2020), is also lower on the X relative to the autosomes (*P* < 0.001, Wilcoxon rank sum test; **Figure 4(b)**). Finally, the density of repeats is higher on the X relative to the autosomes (*P* < 0.001, Wilcoxon rank sum test; **Figure 4(c)**), again consistent with low recombination and/or a higher density of constitutive heterochromatin. **Figure S9** presents these results per scaffold. Interestingly, the nature of repeats appears to differ between the X and the autosomes. While the former seems to have a high proportion of DNA transposons, the autosomes seem to have a higher proportion of retrotransposons (**Table S3**).

**Figure 4:**
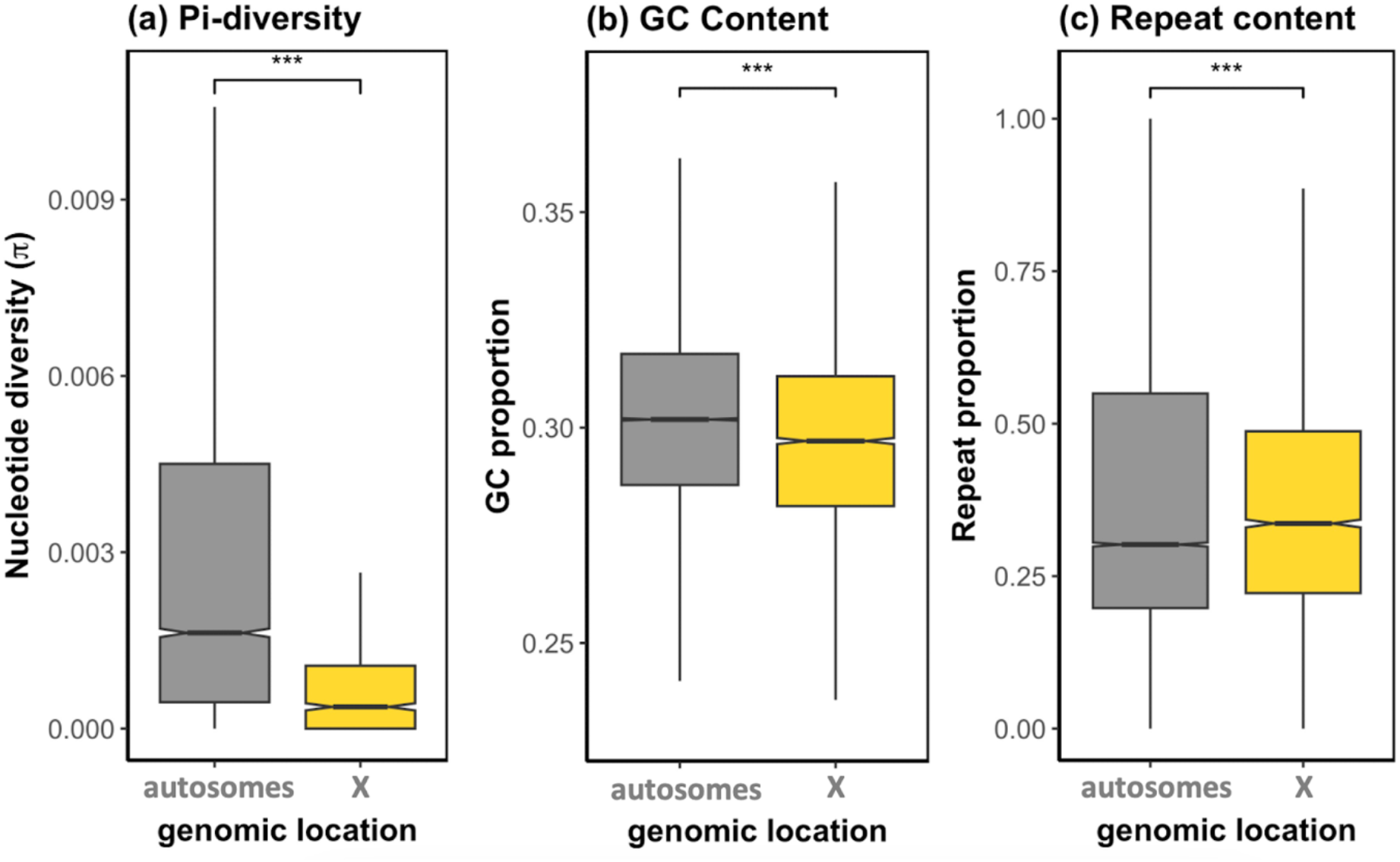
X vs. autosomal (a) female nucleotide diversity (*π*), (b) GC content and (c) repeat content (per 10000 bp windows). Statistically significant differences between groups were assessed using a Wilcoxon rank sum test (* adj. *P* < 0.05, ** adj. *P* < 0.01, *** adj. *P* < 0.001, NS not significant).

## Discussion

### Conservation of the Diptera Muller element F

Our results show that the *P. cognata* X chromosome is homologous to the X of Orthoptera and Blattodea, as well as to the ancient X chromosome of Diptera - Muller element F, consistent with the finding of high conservation of the X chromosome across numerous insect taxa (Meisel et al. 2019; Chauhan et al. 2021; Li et al. 2022; Toups and Vicoso 2023). Despite the homology between the scorpionfly and dipteran X chromosomes, the two chromosomes differ at several key features. First, the dipteran Muller element F is known for its small size; the *P. cognata* X is a much larger chromosome and contains over 1000 genes. This nicely illustrates how “homologous chromosomes” can acquire vastly different gene contents over time due to inter- and intrachromosomal rearrangements, and supports the idea that shrinking of the dipteran X may have driven its high rate of turnover (Toups and Vicoso 2023). Second, our *P. cognata* genome assembly is consistent with a XX/XO male-heterogrametic system, as no scaffolds showed male-specific genomic coverage. The absence of a Y chromosome necessarily implies that sex determination is dosage-dependent, either through the X:autosome ratio - as in Drosophila (Gilbert 2000), or through the number of X chromosomes present in an individual (Blackmon et al. 2017). Because sex-determination is controlled by a Y-linked male-determining factor in some dipterans using the ancestral element F as their X (Sharma et al. 2017; Meccariello et al. 2019; Fan et al. 2023), genes controlling the primary sex determination signal are likely different between Diptera and Mecoptera. This illustrates how the sex determination signal can change even when homology of the sex chromosomes is maintained, and raises the question of what then maintains sex chromosomes over very long periods of time. In mammals, the high conservation of synteny of the X is thought to be driven by the unusual regulatory architecture of this chromosome due to dosage compensation (Ohno 1967; Brashear et al. 2021). A similar argument may apply to insects, since Muller element F is known not only for its specific regulatory mechanisms, but also for being highly heterochromatic and non-recombining.

### The conserved heterochromatic nature of the X

Although characterizing the chromatin and recombinational landscape of the *P. cognata* X would require additional data, we estimated several parameters that are often associated with heterochromatic and/or low recombination regions of the genome: genetic diversity, GC content and repeat content. Similarly to the Diptera Muller element F, the *P. cognata* X appears to be to some extent heterochromatic. In particular, we detected dramatically reduced levels of nucleotide diversity on the X relative to the autosomes (X/A ratio well below 0.75 in both sexes), elevated repeat content on the X relative to the autosomes, and reduced GC content of the X compared to the autosomes. The cockroach X, which is also homologous to element F, is heterochromatic over much of its length (Keil and Ross 1984), raising the possibility that this is an ancestral feature that has contributed to the conservation of this sex chromosome over 450 million years. The characterization of the chromatin landscape of various insects that have maintained the ancestral X chromosome will be needed to shed light on whether its unusual epigenetic profile has played a role in its conservation.

### Partial evidence of demasculinisation of the P. cognata X chromosome

As the X chromosome spends twice as much time in females as in males, sexually antagonistic selection may favour the accumulation of female-beneficial mutations on the X chromosome (Rice 1984; Connallon and Clark 2010). Numerous studies have reported a non-random distribution of genes with sex-biased expression across the genome, with a demasculinisation of the X in numerous taxa, except mammals (Lercher 2003; Zhang et al. 2010). In insects, female-biased genes are generally over-represented on the X relative to the autosomes in Drosophila and beetles (Prince et al. 2010), and reciprocally, male-biased genes seem to escape the X in these two taxa and in mosquitoes (Diptera) (Betrán et al. 2002; Meisel et al. 2009; Vibranovski et al. 2009; Toups and Hahn 2010; Magnusson et al. 2012; Pease and Hahn 2012). Whether the widespread pattern of demasculinisation of the X in insects is a consequence of selection against genes with male-specific functions, or is simply due to reduced expression of the X in testis, is still unclear. The *P. cognata* X shows mixed evidence of demasculinisation of the X. On the one hand, genes with male-biased expression appeared less prevalent on the X than on the autosomes, but, on the other hand, genes exclusively expressed in testis were equally as common on the X and the autosomes (**Figure 3(e) and (f)**). Our results therefore show that genes that function primarily in the testes can survive on the X even when the expression of this chromosome is generally female-biased, perhaps arguing against the hypothesis that selection has driven male-biased genes out of the X.

## Data accessibility

All raw RNA-seq and DNA-seq data have been uploaded to the NCBI under project PRJNA989034. Processed data files are available at: https://seafile.ist.ac.at/d/efa3989c33024b859c02/. Pipelines are available at: https://github.com/ClemLasne/PanorpaX

## Supplementary information

Please see Supplementary_material_Panorpa_manuscript.pdf

## Authors’ contribution

CL and ME: conceptualization, data curation, formal analysis, methodology, writing—original draft, writing—review and editing; MT and LL: formal analysis, writing—original draft, writing—review and editing; A.M.: methodology, resources; B.V.: conceptualization, formal analysis, funding acquisition, project administration, writing—original draft, writing—review and editing. All authors gave final approval for publication and agreed to be held accountable for the work performed therein.

## Acknowledgements

We thank the Vicoso lab for their assistance with specimen collection, and Tim Connallon for valuable comments and suggestions on earlier versions of the manuscript. Computational resources and support were provided by the Scientific Computing unit at the ISTA. This research was supported by grants from the Austrian Science Foundation to C.L. (FWF ESP 39), and to B.V. (FWF SFB F88-10).

## Notes

### Competing Interest Statement

The authors have declared no competing interest.

