## Supplementary material for "The scorpionfly (*Panorpa cognata*) genome highlights conserved and derived features of the peculiar dipteran X chromosome"


#### SUPPLEMENTARY DATASETS 2

**Dataset S1: Genome assembly 2**

**Dataset S2. Transcriptome assembly and transcripts genomic location 2**

**Dataset S3: Identification of X-linked scaffolds 2**

**Dataset S4: Gene expression and dosage compensation 2**

#### SUPPLEMENTARY METHODS 3

**Methods S1: Homology of *Drosophila melanogaster* to *Panorpa cognata* 3**

**Methods S2: X-linked genes in Blatella germanica and homology to *Panorpa cognata* 3**

#### SUPPLEMENTARY FIGURES 4

**Figure S1: BUSCO assessments of our genome and transcriptome assemblies. 4**

**Figure S2: Male and female genome coverage for all scaffolds. 4**

**Figure S3: Homology of the X chromosomes of *P. cognata* (order: Mecoptera) and three other insects: two Diptera, *C. hominivorax* (suborder: Brachycera) and *N. appendiculata* (suborder: Nematocera), and *L. migratoria* (order: Orthoptera). 6**

**Figure S4: Percentage of genes on each of the *D. melanogaster* Muller elements that are X-linked in *P. cognata*. 6**

**Figure S5: Percentage of X-linked and autosomal *P. cognata* genes that are X-linked in *B. germanica* (order: Blattodea). 7**

**Figure S6: Spearman correlation heatmap displaying the overall similarity in expression profiles of the RNA-seq samples. 8**

**Figure S7: Dosage compensation in carcasses. 8**

**Figure S8: Expression level of gene *Painting Of Fourth* (*POF*) in each RNA-seq sample. 9**

**Figure S9: Detailed per-scaffold nucleotide diversity, GC and repeat content. 10**

#### SUPPLEMENTARY TABLES 11

**Table S1: List of RNA and DNA samples generated and the analysis they were used for. 11**

**Table S2: Statistics for the different assembly steps of the *Panorpa cognata* genome. 11**

**Table S3: The classification and distribution of repetitive content on the autosomes and the X chromosome. 11**

### Supplementary Datasets

**Dataset S1: Genome assembly**
**Dataset S2. Transcriptome assembly and transcripts genomic location**
**Dataset S3: Identification of X-linked scaffolds**
**Dataset S4: Gene expression and dosage compensation**
**Dataset S5: GC content and nucleotide diversity**
**Dataset S6: Repeat Content**
**Dataset S7: Homology (and orthofinder)**

**All supplementary datasets are available at:**
https://seafile.ist.ac.at/d/efa3989c33024b859c02/

### Supplementary Methods

##### Methods S1: Homology of *Drosophila melanogaster* to *Panorpa cognata*

*Drosophila melanogaster* protein sequences were retrieved from Ensembl (https://ftp.ensembl.org/pub/release-108/fasta/drosophila_melanogaster/pep/). We filtered for the longest sequence of each protein using a Perl script (GetLongestCDS_v2.pl). We used the *D. melanogaster* gene names and their Muller element location according to the CDS from (https://ftp.ensembl.org/pub/release-108/fasta/drosophila_melanogaster/cds/). We used Orthofinder (Emms and Kelly 2019) to obtain the 1-to1 orthologs of *D. melanogaster* and *P. cognata* using the same pipeline as explained in our paper in the "***Homology of the Panorpa cognata and Cochliomyia hominivorax X chromosomes***" methods section.

##### Methods S2: X-linked genes in *Blatella germanica* and homology to *Panorpa cognata*

We used published *Blatella germanica* DNA reads from two sources: (1) male reads (SRR1566154, SRR1566155, SRR1566159) and one female reads (SRR1566152) from Meisel et al. (2019); and (2) two male reads (SRR9160163, SRR9160164) from NCBI bioproject number PRJNA545466. The female reads from the same bioproject seemed incorrectly labelled and therefore were not used in our analysis. We mapped the reads to the *Blatella germanica* genome (Meisel et al. 2019) using Bowtie2 (version 2/2.4.5 (Langmead and Salzberg 2012)) and then we used SOAP.coverage (version 2.7.7; https://github.com/gigascience/bgi-soap2/tree/mas-ter/tools/soap.coverage) to calculate the coverage depth. As we had five male reads in total, their coverage depth was summed up. We only kept the scaffolds with length greater than 1000 bp. Finally, we calculated the $\log_2$ of the ratio of male to female coverage ($Log_2$(Male/Female coverage)) for each scaffold. The scaffolds were assigned as X-linked if that ratio was less than the [median($Log_2$(Male/Female coverage))-0.5], otherwise they were assigned as autosomal. In order to assess the homology of *B. germanica* and *P. cognata*, we used the same pipeline as explained in our paper in the "***Conservation of X-linked gene content between P. cognata and other insects***" methods section.

### Supplementary Figures

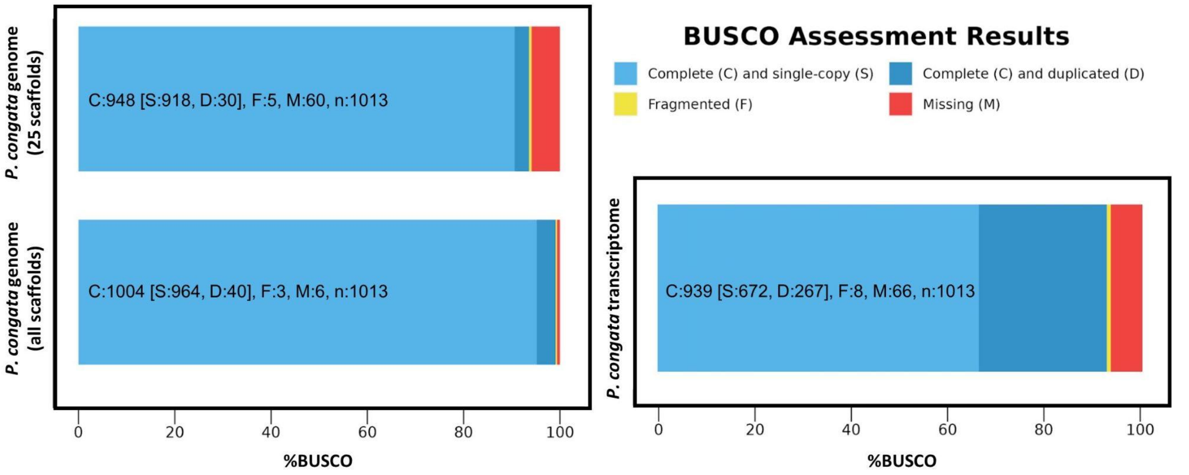


**Figure S1: BUSCO assessments of our genome and transcriptome assemblies.**

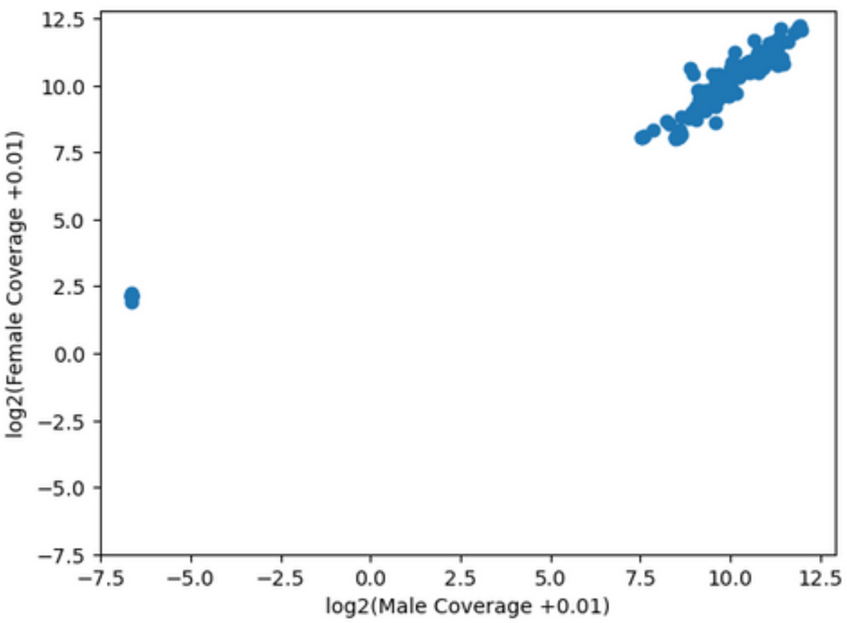


**Figure S2: Male and female genome coverage for all scaffolds.**

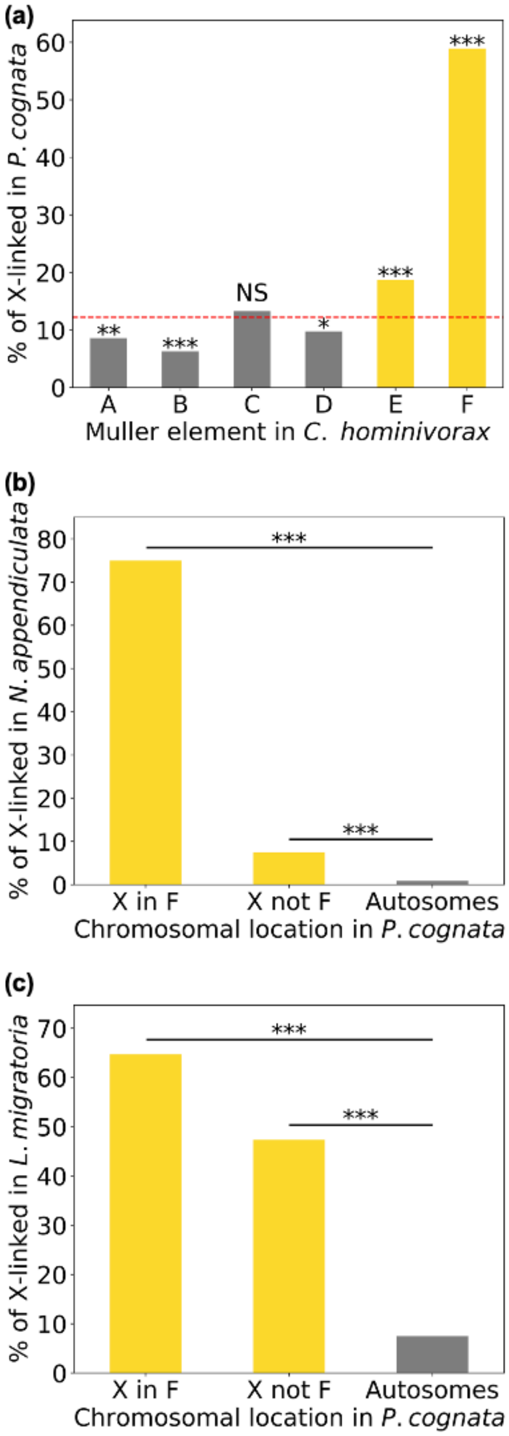
(a)
% of X-linked in P. cognata
60
50
40
30
20
10
0
***
***
NS
**
***
*
A
B
C
D
E
F
Muller element in C. hominivorax
(b)
% of X-linked in N. appendiculata
80
70
60
50
40
30
20
10
0
***
***
X in F
X not F
Autosomes
Chromosomal location in P. cognata
(c)
% of X-linked in L. migratoria
70
60
50
40
30
20
10
0
***
***
X in F
X not F
Autosomes
Chromosomal location in P. cognata

**Figure S3: Homology of the X chromosomes of *P. cognata* (order: Mecoptera) and three other insects: two Diptera, *C. hominivorax* (suborder: Brachycera) and *N. appendiculata* (suborder: Nematocera), and *L. migratoria* (order: Orthoptera).** All the scaffolds in our P. cognata genome were taken into account. (a) Percentage of genes on each of the C. hominivorax's Muller elements that are X-linked in P. cognata. The red dashed line represents the overall proportion of orthologs that are X-linked in P. cognata (i.e. the "X-linkage threshold"). (b) Percentage of X-linked and autosomal P. cognata genes that are X-linked in N. appendiculata. The X-linked genes of P. cognata were divided into two sets, based on whether they were F-linked in C.hominivorax (X-in-F), or not (X-not-F). (c) same as (b) but showing the percentage of P. cognata genes that are X-linked in L. migratoria. Statistically significant differences between observed and expected percentages were assessed using a chi-squared test (* P $<$ 0.05, ** P $<$ 0.01, *** P $<$ 0.001, NS not significant).

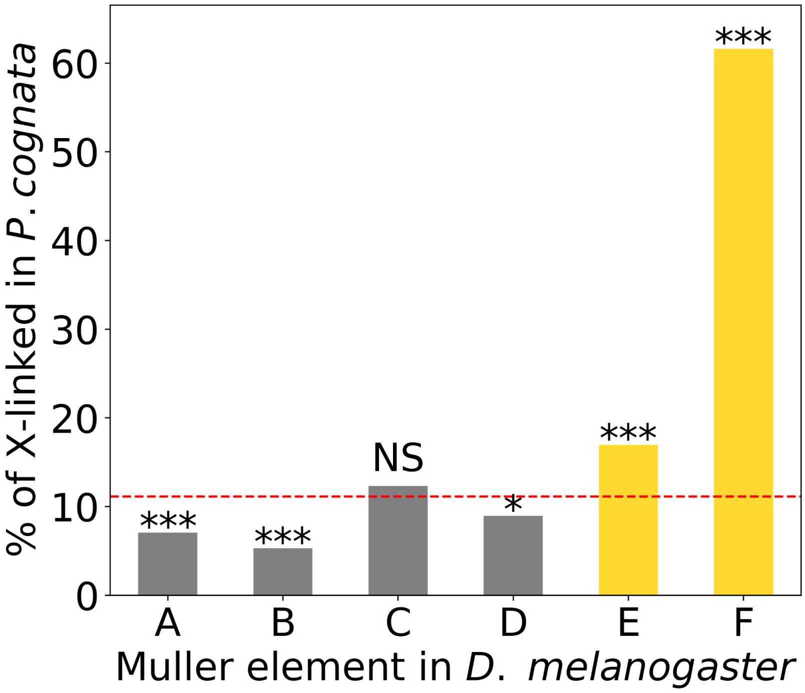


**Figure S4: Percentage of genes on each of the *D. melanogaster* Muller elements that are X-linked in *P. cognata.*** This is based on 1-to-1 orthologs analysis and the *P. cognata*'s 25 largest genome scaffolds. The red dashed line represents the overall proportion of orthologs that are X-linked in *P. cognata* (i.e. the "X-linkage threshold"). Statistically significant differences between observed and expected percentages were assessed using a chi-squared test (* $P < 0.05$, ** $P < 0.01$, *** $P < 0.001$, NS not significant).

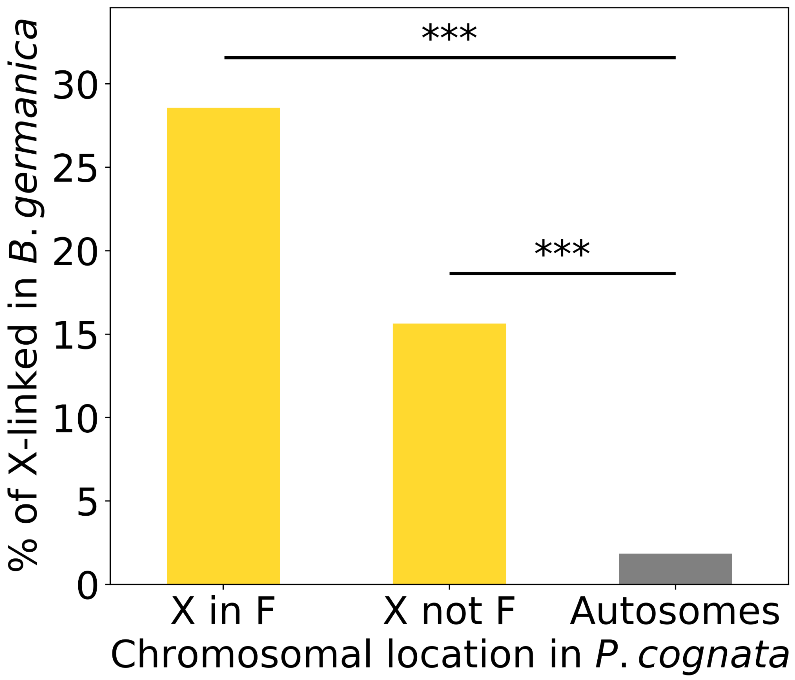


**Figure S5: Percentage of X-linked and autosomal *P. cognata* genes that are X-linked in *B. germanica* (order: Blattodea).** The X-linked genes of *P. cognata* were divided into two sets, based on whether they were F-linked in *C.hominivorax* (X-in-F), or not (X-not-F). Statistically significant differences between observed and expected percentages were assessed using a chi-squared test (*** $P < 0.001$).

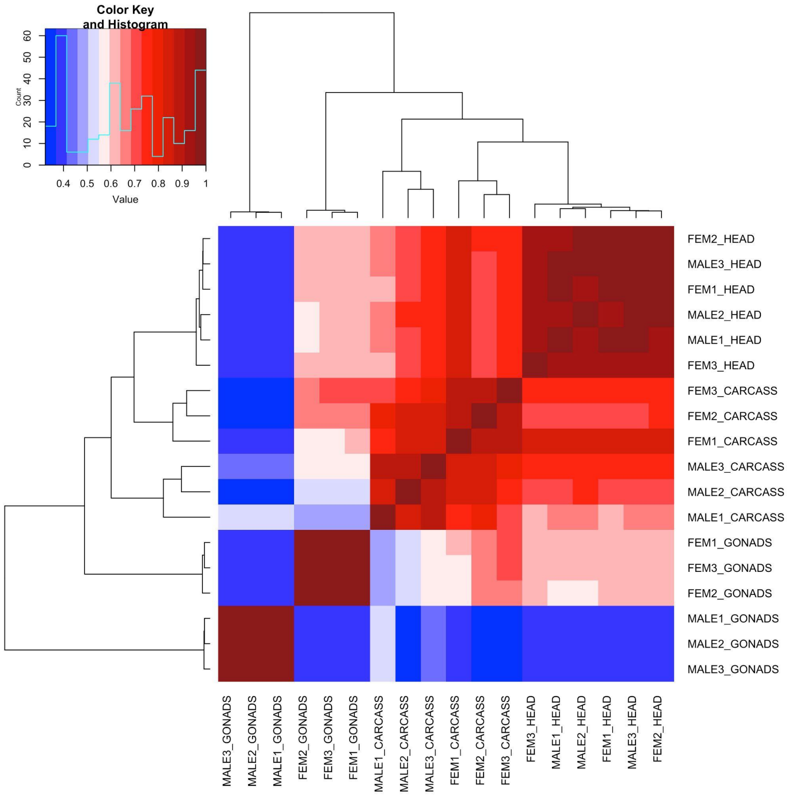

**Figure S6: Spearman correlation heatmap displaying the overall similarity in expression profiles of the RNA-seq samples.** This is based on the 13785 genes of known chromosomal location. Sample IDs are indicated on the x-axis and secondary y-axis. Similarity between samples is indicated by the dendrogram on the y-axis and the secondary x-axis. Correlation intensity between samples increases from blue (low correlation) to red (high correlation).

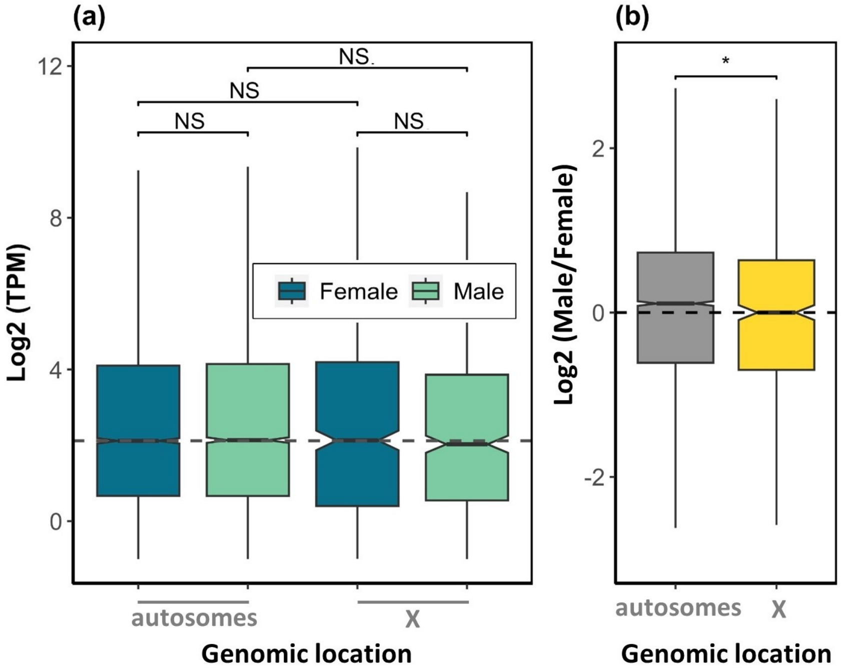


**Figure S7: Dosage compensation in carcasses.** (a) Expression of autosomal and X-linked genes in males and females (grey dashed line is the female autosomal gene expression median). (b) $Log_2$ of male-over-female expression ratios for the autosomal and X-linked genes. Statistically significant differences between groups were assessed using a Wilcoxon rank sum test (* $P < 0.05$, ** $P < 0.01$, *** $P < 0.001$, NS not significant).

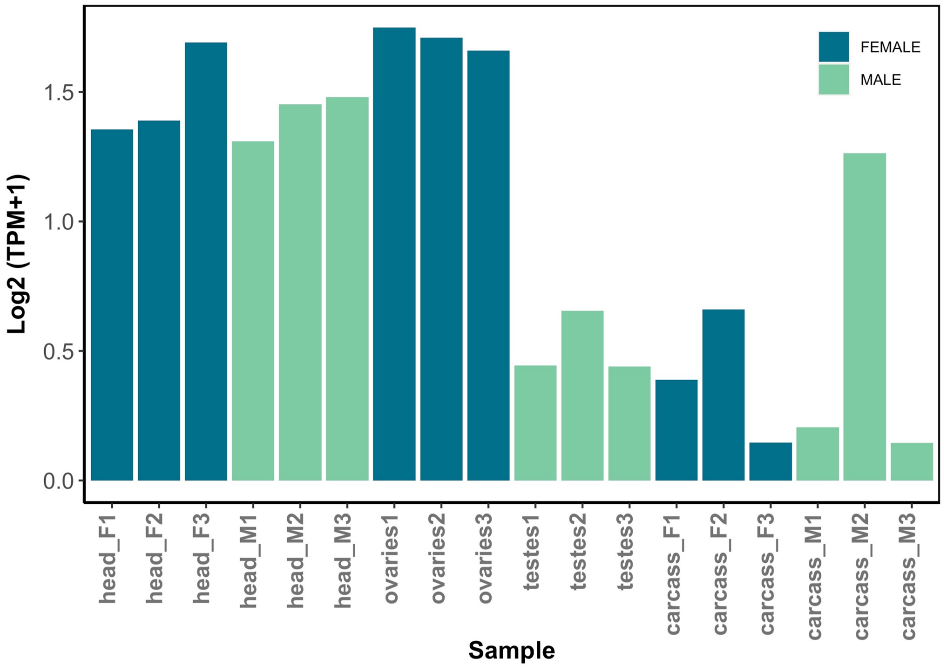


**Figure S8: Expression level of gene *Painting Of Fourth* (*POF*) in each RNA-seq sample.**

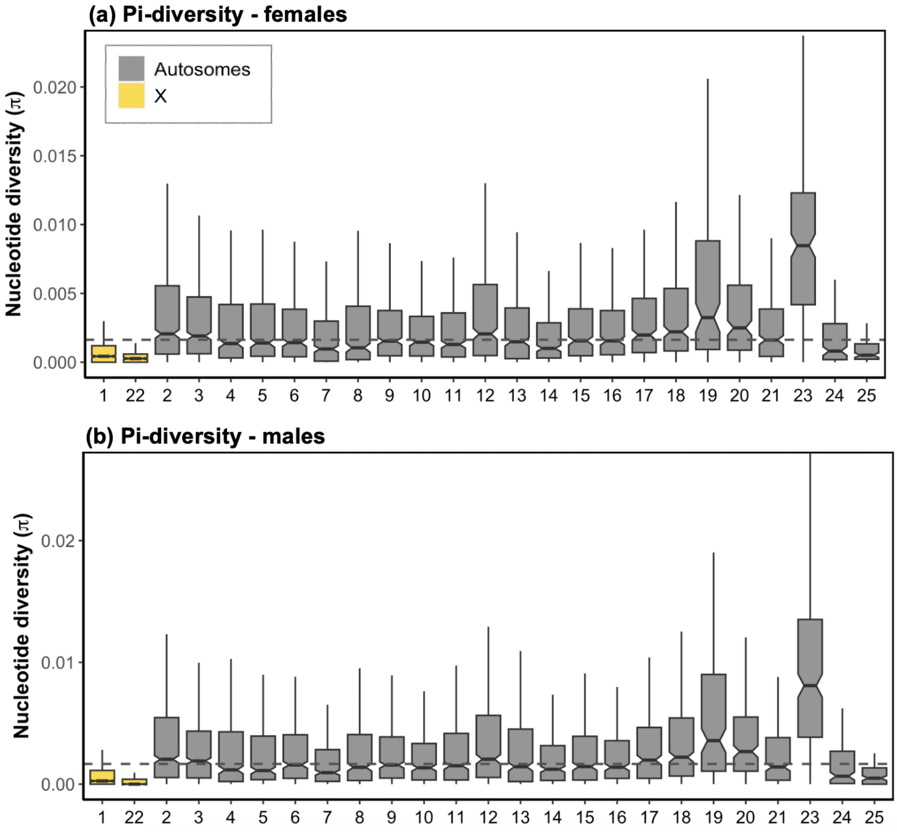

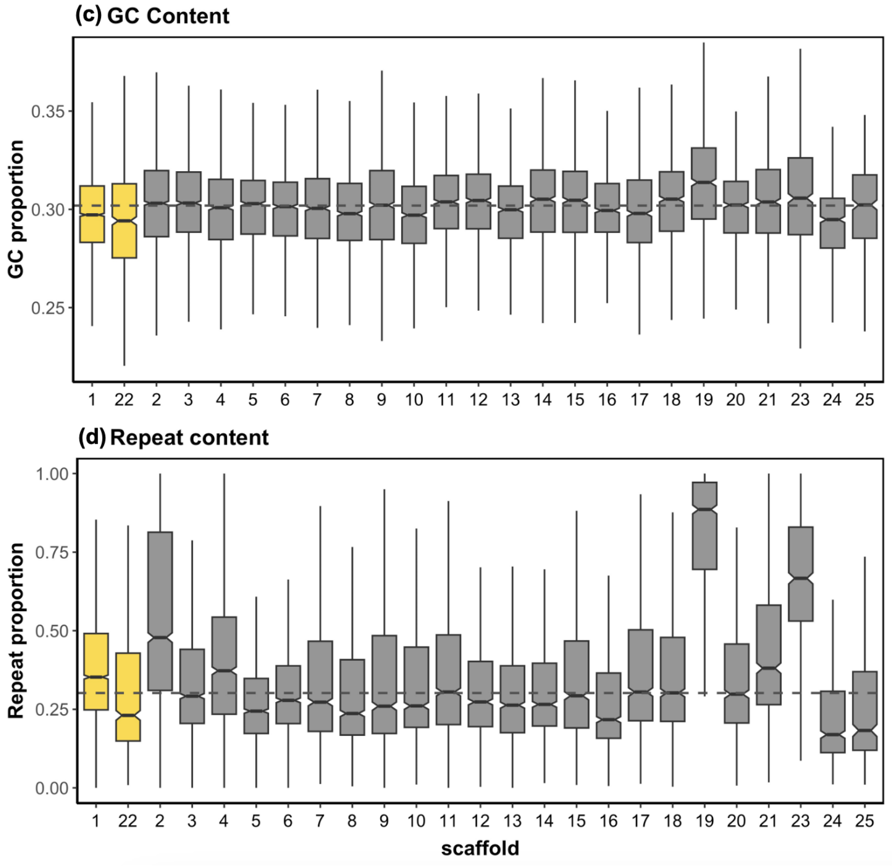


**Figure S9: Detailed per-scaffold nucleotide diversity, GC and repeat content.** (a and b) female and male nucleotide diversity, (c) GC content (per 10000 bp windows) and (d) repeat content (per 10000 bp windows).

### Supplementary Tables

**Table S1: List of RNA and DNA samples generated and the analysis they were used for.**

Available at : [https://seafile.ist.ac.at/library/b0536613-0190-4733-9f8d-48863df51131/panorpaX/](https://seafile.ist.ac.at/library/b0536613-0190-4733-9f8d-48863df51131/panorpaX/)

**Table S2: Statistics for the different assembly steps of the *Panorpa cognata* genome.**

| | hifiasm | purged | Hi-C scaffolded (scaffolds longer >N80) | Longest 25 scaffolds |
|---|---|---|---|---|
| size | 1034432113 | 571169746 | 457046408 | 333218704 |
| n | 6056 | 1359 | 187 | 25 |
| N50 | 490137 | 9373981 | 11774435 | 13885755 |
| N90 | 57472 | 137127 | 616955 | 9075028 |
| largest | 18224189 | 18224189 | 35185269 | 35185269 |
| Average | 170811.12 | 420286.79 | 2444098.44 | 13328748.16 |
| N_count | 0 | 0 | 2200 | 1800 |
| Gaps | 0 | 0 | 11 | 9 |

**Table S3: The classification and distribution of repetitive content on the autosomes and the X chromosome.**

| | Autosomes (23 scaffolds) | | | X chromosome (2 scaffolds) | | |
|---|---|---|---|---|---|---|
| | #Elements | Length occupied (bp) | % sequence | #Elements | Length occupied (bp) | % sequence |
| Retroelements | 34008 | 16659493 | 5.74 | 4308 | 1848169 | 4.28 |
| DNA transposons | 47485 | 10109131 | 3.49 | 8945 | 1844182 | 4.27 |
| Rolling-circles | 6381 | 1237671 | 0.43 | 1216 | 282311 | 0.65 |
| Unclassified | 440128 | 79688795 | 27.48 | 70177 | 11944909 | 27.64 |
| Small RNA | 11969 | 3445476 | 1.19 | 1953 | 619645 | 1.43 |

| | | | | | | |
|---|---|---|---|---|---|---|
| Satellites | 338 | 147685 | 0.05 | 18 | 1729 | 0 |
| Simple repeats | 96244 | 5095369 | 1.76 | 17296 | 952287 | 2.2 |
| Low complexity | 17721 | 859362 | 0.3 | 2804 | 136626 | 0.32 |
